## Supplementary material for "N-myc Downstream Regulated Gene 1 (NDRG1) functions as a molecular switch for cellular adaptation to hypoxia": Source Data Legends

Figure 1-figure supplement 1-Source Data 1. Source data of uncropped, annotated WB figures of Ndrg1a and GAPDH; Raw, unedited WB file for Ndrg1a; Raw, unedited WB file for GAPDH

Figure 1-figure supplement 3-Source Data 1. Source data for quantification of kidney clearance assay under normoxia (48 hpf) WT and *ndrg1a*^-/-^ mutant.

Figure 2-Source Data 1. Source data for % survival of WT embryos and *ndrg1a^-/-^* mutants immediately following 6, 12, 18 and 24 h of anoxia exposure.

Figure 2-Source Data 2. Source data for % survival of WT embryos and *ndrg1a*^-/-^ mutants following 6, 12, 18 and 24 h of anoxia exposure and 2 days of re-oxygenation (re-ox).

Figure 2-Source Data 3. Source data for % edema observed in WT embryos and *ndrg1a*^-/-^ mutants following 0, 6, 12, 18 and 24 h of anoxia exposure and 2 days re-ox.

Figure 2-Source Data 4. Source data for kidney clearance assay using 48 hpf WT embryos and *ndrg1a*^-/-^ mutants exposed to 12 h of anoxia, followed by 2 days re-ox and injection with rhodamine dextran.

Figure 3-Source Data 1. Source data for normalized fluorescence intensity in the (c1) anterior and (c2) posterior pronephric duct of WT embryos and *ndrg1a^-/-^* mutants.

Figure 3-figure supplement 1-Source Data 1. Source data for quantification of ATP1A1A levels during reoxygenation.

Figure 4-Source Data 1. Source data for normalized fluorescence intensity of ATP1A1A in ionocytes of WT embryos and *ndrg1a^-/-^* mutants exposed to anoxia for 0, 6, 12, 18, and 24h.

Figure 5-Source Data 1. Source data for quantification of ATP1A1A fluorescence intensity in the pronephric duct of embryos subjected to 0 h (d1) or 12 h of anoxia (d2-d5) in presence or absence of MG-132, or chloroquine, or both and immunolabeled using anti-ATP1A1A.

Figure 5-Source Data 2. Source data for quantification of total ATP concentration per WT embryo or *ndrg1a^-/-^* mutant exposed to 0, 6, 12 h of anoxia.

Figure 5-Source Data 3. Source data for quantification of total ATP concentration per WT embryo exposed to 12h of anoxia with or without ouabain.

Figure 5-figure supplement 1-Source Data 1. Source data for ITC experiments.

Figure 6-Source Data 1. Source data for Figure 6A: Metabolites enriched in anoxia-treated embryos relative to normoxic controls; Figure 6B: Extracted ion counts for different elution timepoints are shown for two extracts from sphere stage (4 hpf) embryos exposed to 1 h of anoxia (orange and red lines) and two control samples from 5 hpf normoxic embryos (yellow and green lines); Figure 6C: Comparison of lactate levels (highest peak intensity) in extracts from sphere stage (4 hpf) embryo exposed to 1 h of anoxia and 5 hpf normoxic controls reveals an 18 fold increase.

Figure 6-Source Data 2. Source data for fluorometric lactate assay showing lactate concentration (nmol lactate/embryo) in whole embryo extracts from 24 hpf WT or *ndrg1a^-/-^* mutant exposed to 0, 1, 3, 6, 12 h of anoxia.

Figure 7-Source Data 1. Source data for fluorometric lactate assay showing lactate concentration (nmol lactate/embryo) in whole embryo extracts from 24 hpf WT or *ndrg1a^-/-^* mutant exposed to sodium azide for 0, 3, 6, 12 hours under normoxia.

Figure 7-Source Data 2. Source data for normalized NKA fluorescence intensity in the anterior (circle) and posterior (triangle) pronephric duct of WT (black) embryos and *ndrg1a^-/-^* (red) mutants.

Figure 7-Source Data 3. Source data for normalized PLA intensity in the anterior (circle) and posterior (triangle) pronephric duct of WT embryo.
